# Semi-supervised sequence modeling for improved behavioral segmentation

**DOI:** 10.1101/2021.06.16.448685

**Authors:** Matthew R Whiteway, Evan S Schaffer, Anqi Wu, E Kelly Buchanan, Omer F Onder, Neeli Mishra, Liam Paninski

**Author notes:** An abbreviated version of this work is to appear at the CVPR 2021 CV4Animals Workshop.

## Abstract

A popular approach to quantifying animal behavior from video data is through discrete behavioral segmentation, wherein video frames are labeled as containing one or more behavior classes such as walking or grooming. Sequence models learn to map behavioral features extracted from video frames to discrete behaviors, and both supervised and unsupervised methods are common. However, each approach has its drawbacks: supervised models require a time-consuming annotation step where humans must hand label the desired behaviors; unsupervised models may fail to accurately segment particular behaviors of interest. We introduce a semi-supervised approach that addresses these challenges by constructing a sequence model loss function with (1) a standard supervised loss that classifies a sparse set of hand labels; (2) a weakly supervised loss that classifies a set of easy-to-compute heuristic labels; and (3) a self-supervised loss that predicts the evolution of the behavioral features. With this approach, we show that a large number of unlabeled frames can improve supervised segmentation in the regime of sparse hand labels and also show that a small number of hand labeled frames can increase the precision of unsupervised segmentation.

## 1. Introduction

Behavioral segmentation is an indispensable tool for quantifying natural animal behavior as well as understanding the effects of targeted interventions [1, 35, 44]. This procedure begins with the collection of raw behavioral data during an experiment, typically with video or motion-capture sensors. In the supervised segmentation setting, the experimenter then labels a subset of frames that contain behaviors of interest, such as walking, grooming, rearing, etc. Finally, a machine learning algorithm (referred to here as a “sequence model”) is trained to match each frame with the corresponding behaviors [16, 18, 20, 31, 32, 6, 33, 39, 41, 43]. As the scale of behavioral data continues to grow [11, 44], it becomes infeasible to densely label behaviors in every video. Therefore, it is crucial to develop segmentation techniques that perform well with sparsely labeled data.

Fully unsupervised behavioral segmentation algorithms are a complementary approach that require no hand labels [4, 17, 47, 14, 26]. These algorithms typically reduce the dimensionality of the raw video data through various methods, then perform unsupervised clustering on the resulting low-dimensional behavioral representation [9]. These unsupervised methods tend to be more scalable than their supervised counterparts because they do not require manual input. Another benefit of these methods is their ability to discover new behaviors [9, 35]. However, there may be certain behaviors of particular interest for downstream analyses, and unsupervised methods cannot guarantee they accurately segment these behaviors.

Here we propose a semi-supervised segmentation algorithm that combines the strengths of these two approaches while minimizing their weaknesses: we take advantage of a small number of hand labels to ensure particular behaviors are well-represented by the model and also take advantage of a large number of unlabeled frames in order to improve the behavioral representation (Fig. 1). We find that this approach improves supervised segmentation while still allowing for unsupervised behavior discovery.

**Figure 1.**
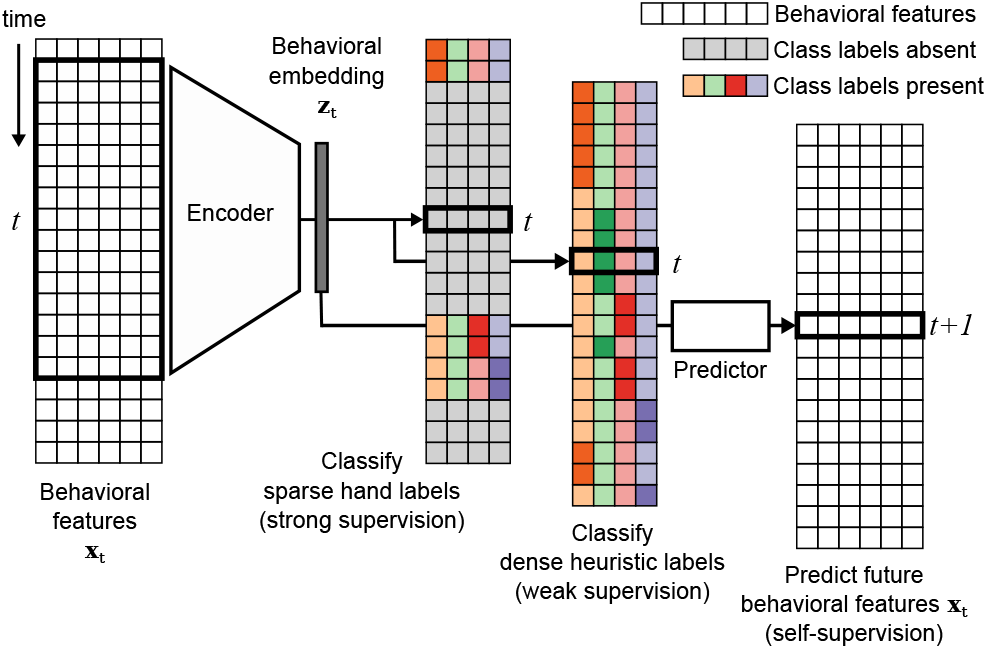
Temporal sequence models for behavioral classification can be augmented with a weakly supervised loss that classifies heuristic labels and a self-supervised loss that predicts future behavioral features in order to improve model performance on sparsely labeled datasets.

We propose two simple losses that extract relevant information from unlabeled frames, which augment a standard supervised classification loss computed on labeled frames (e.g. cross entropy). The first loss is based on the observation that the behaviors of interest can often be described heuristically: for example, when tracking the paws and the nose of a mouse with pose estimation software [29], candidate bouts of grooming can be identified by finding all frames where the distance from the paws to the nose is below some threshold. This procedure may lead to false positives (e.g. when the mouse grabs a lick spout near its nose) and false negatives (when the distance threshold is set too low), but it is a straightforward and computationally efficient way to automatically label many frames in a video. We propose, then, to use these so-called “heuristic” labels as an additional supervised signal (which we refer to as *weak* supervision) for sequence models.

The second loss is based on the observation that behavior is *dynamic*; even a “still” behavior is defined in the context of what the animal was doing before and after a “still” frame. Therefore, we propose to add a self-supervised prediction loss so the model learns to map the behavioral features at time *t* to the behavioral features at time *t*+1. Note that this prediction loss can be computed for every single frame of a video, without the need for corresponding hand or heuristic labels, again allowing the sequence model to take advantage of potentially large amounts of unlabeled data.

We evaluate our proposed semi-supervised approach by conducting an empirical evaluation on a head-fixed fly dataset. We find that, individually, the weak and self-supervised losses improve supervised segmentation metrics across all behaviors; combining these losses leads to additional improvements. We also compare our approach to fully unsupervised behavioral segmentation models and show that adding a small number of hand labels can improve segmentation while still allowing for the discovery of previously unlabeled behaviors. Code is available at https://github.com/themattinthehatt/daart.

### Related Work

We focus on behavioral segmentation using pose estimates, and therefore our work draws from the literature on skeleton-based action understanding [13]. [10] shows that motion prediction is a good auxiliary task for behavioral segmentation when using sparse hand labels; we build upon this work by including an additional set of heuristic labels that can strengthen the classifier even further. [42] introduces a set of “task program” heuristics to shape a latent behavioral embedding used by downstream behavior classifiers; while this is similar in spirit to our approach, we choose instead to provide direct heuristics for each hand-labeled behavior, and train our model end-to-end. [24, 25] use a classification and prediction loss to perform semi-supervised action recognition (different from segmentation), though their focus is on an active learning scheme for determining the most informative data points for future labeling. Several other works introduce weak supervision terms to improve segmentation [7, 15, 21, 36], though they rely on relatively strong assumptions, such as a known ordering of behaviors, that are not relevant for our use case.

If we use our model with the self-supervised loss only, we can perform fully unsupervised behavioral segmentation by performing a post-hoc clustering of the low-dimensional behavioral embeddings. This approach follows a common pipeline successfully used across many studies [9]. For example, [26] use an autoencoder RNN to produce a behavioral embedding from pose estimates, apply UMAP [30] to further reduce the dimensionality, then apply k-means clustering to perform unsupervised behavioral segmentation. Other recent works use different combinations of algorithms for embedding, dimensionality reduction, and clustering [4, 47, 17, 3, 27]. Our work expands this pipeline to include hand and heuristic labels, and we show that this semi-supervised approach can produce higher quality segmentations.

## 2. Methods

Most approaches for supervised behavioral segmentation from video data involve a two-step process: first, for each frame **f**_*t*_, compute a lower-dimensional feature representation **x**_*t*_ that encodes local spatiotemporal information using pose estimation [28, 12, 34, 48], Dense Trajectories [45], or two-stream network outputs [40, 6]; second, a sequence model *f* (·) (such as a recurrent neural network) maps the behavioral feature vector **x**_*t*_ (or a window of these features) to a discrete label vector **ŷ**_*t*_, which should match the hand labels **y**_*t*_. We assume that the hand labels are only defined on a subset of time points 𝒯 ⊆ {1, 2, …*T*}. The crossentropy loss function ℒ_xent_ [5] then defines the supervised objective (ℒ_super_) to optimize:

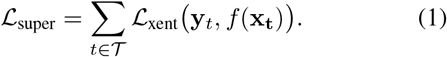

### Weak and self-supervised loss functions

We now introduce a set of heuristic labels 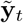, defined at each time point. Computing the cross-entropy loss on all time points that do not already have a corresponding hand label defines the heuristic objective:

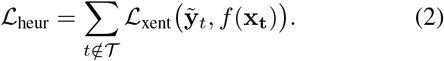

The self-supervised loss requires the sequence model to predict **x**_*t*+1_ from **x**_*t*_. To properly describe this we now expand the definition of the sequence model *f* (·) to include two components: an *encoder e*(·), which maps the behavioral features **x**_*t*_ to an intermediate behavioral embedding **z**_*t*_; and a (linear) *classifier c*(·) which maps **z**_*t*_ to the predicted discrete labels (**ŷ**_*t*_ = *c*(*e*(**x**_*t*_)). We can now incorporate the self-supervised loss through the use of a *predictor* function *p*(·), which maps **z**_*t*_ to 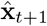, and match 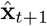 to the true behavioral features **x**_*t*+1_ through a mean square error loss ℒ_MSE_ computed on all time points:

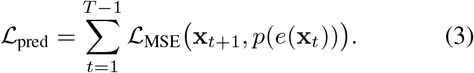

Finally, we combine all terms into the full semi-supervised loss function:

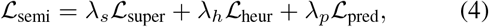

where the *λ* terms are hyperparameters that control the contributions of their respective losses. Note that setting *λ*_*h*_ = *λ*_*p*_ = 0 results in a fully supervised model, while *λ*_*s*_ = *λ*_*h*_ = 0 results in a fully unsupervised model.

### Model

Our approach is agnostic to the particular architecture of the encoder and predictor networks. Here we use a standard recurrent neural network with GRU layers [8], which sees frequent use in sequence modeling [2]. Both *e*(·) and *p*(·) are modeled with two layer bidirectional GRU networks (32 cells per layer). We performed a small hyperparameter search across number of layers, cells per layer, and learning rate, and found that our results are robust across different settings (data not shown).

### Data

We evaluate our approach on a head-fixed fly dataset [38] (Fig. 2). This dataset contains videos from 10 flies, with videos ranging in length from 10 to 34 minutes. We first track eight points on the fly using Deep Graph Pose [48], and use these points as our behavioral features **x**_*t*_ (see Fig. 2). The flies exhibit a small range of easily identified behaviors, including standing still, walking on an air-supported ball, and front and back grooming (Fig. 2, top). We label up to 300 frames for each of these behaviors per fly (for a total of 1.1% of all frames labeled; see Table S1). We also devise a simple set of heuristics to produce weak labels for each behavior: frames are labeled “Walk” when the ball motion energy (ME) is above a threshold; “Still” when the ME of the limb markers is below a threshold; and “Front (Back) groom” when the fly is not walking, and the ME of the forelimb (hindlimb) markers is above a threshold.

**Figure 2.**
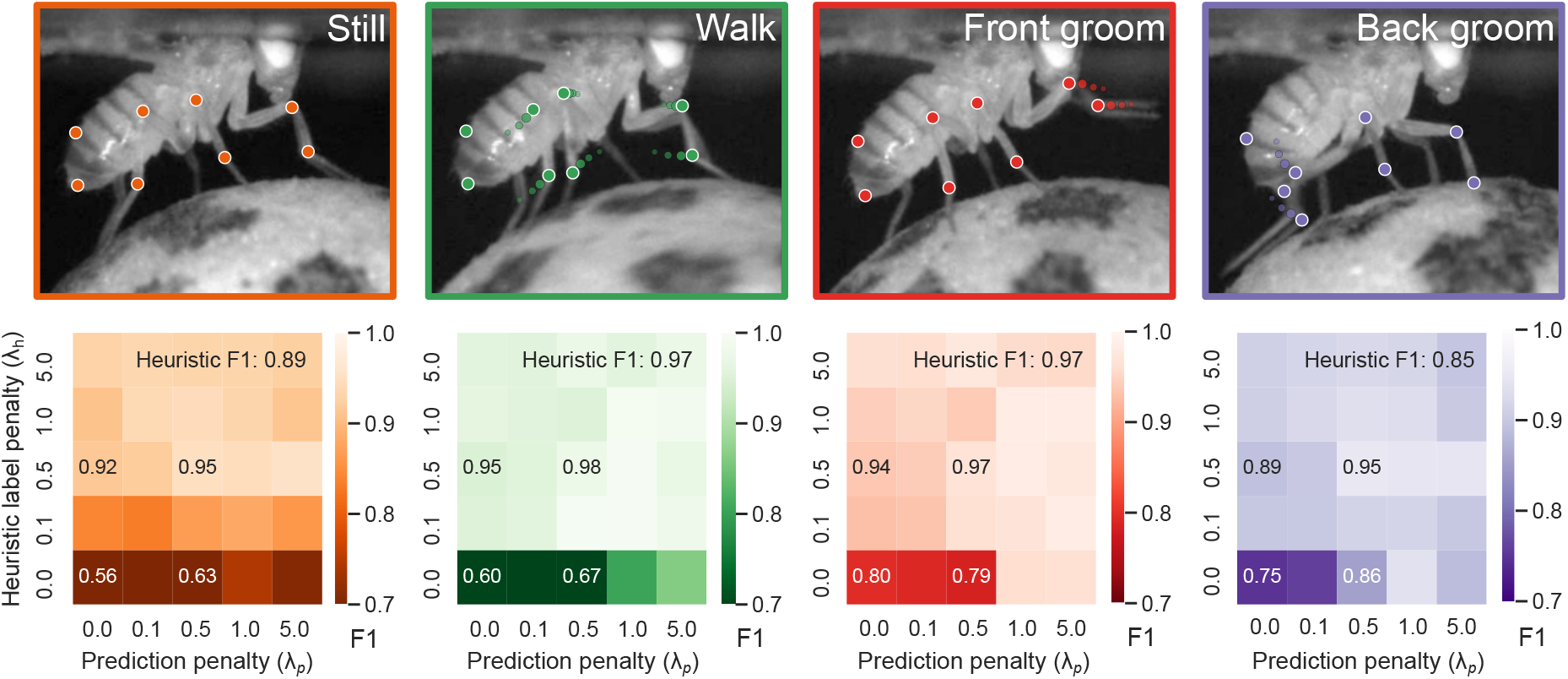
*Top*: Example frames from four fly behavior classes, each overlaid with the eight points tracked through pose estimation. Smaller trailing dots indicate the dynamics of each tracked point during the specified behavior. *Bottom*: F1 score for each behavior on test data as a function of the weak supervised penalty based on heuristic labels (*λ*_*h*_) and the self-supervised prediction penalty (*λ*_*p*_). F1 score is overlaid in text for selected hyperparameter combinations; see Fig. S2 for all values. Higher F1 scores are better, with a maximum of value of 1. Standard supervised classification corresponds to *λ*_*h*_ = *λ*_*p*_ = 0.

### Training and evaluation

We use 5 fly videos for training and 5 for testing. All models are trained with the Adam optimizer [19] using an initial learning rate of 1e− 4 and a batch size of 2000 time points. For the 5 training flies, 90% of frames are used for training, 10% for validation. Training is terminated once the loss on validation data begins to rise for 20 consecutive epochs; the epoch with the lowest validation loss is used for testing. To evaluate the models, we compute the F1 score – the geometric mean of precision and accuracy – on the hand labels of the 5 held-out test flies.

## 3. Results

We first consider the effect of the weak and self-supervised losses on supervised segmentation (and set *λ*_*s*_ = 1). The weak supervision provided by the heuristic labels, controlled by *λ*_*h*_, dramatically improves F1 scores across all four behavior classes, compared to the supervised baseline of *λ*_*h*_ = *λ*_*p*_ = 0 (Fig. 2, bottom). The presence of the high-quality hand labels also allows the model to surpass the F1 score of the lower-quality heuristic labels, especially on the more difficult behaviors (see text overlaid on heatmaps in Fig. 2). Next we consider the effect of self-supervision through the next-step-ahead prediction, controlled by *λ*_*p*_. Nonzero values of *λ*_*p*_ also improve F1 scores across all four behavior classes. Finally, we find that combining both loss functions can slightly increase F1 yet again (for example see F1 scores for *λ*_*h*_ = *λ*_*p*_ = 0.5). Fig. 3 shows marker traces and their behavior class probabilities from the best performing model for several sample time segments; Fig. S1 shows the corresponding class probabilities from the fully supervised model (*λ*_*h*_ = *λ*_*p*_ = 0), and demonstrates how the absence of the weak and self-supervised losses leads to more errors in the segmentation.

**Figure 3.**
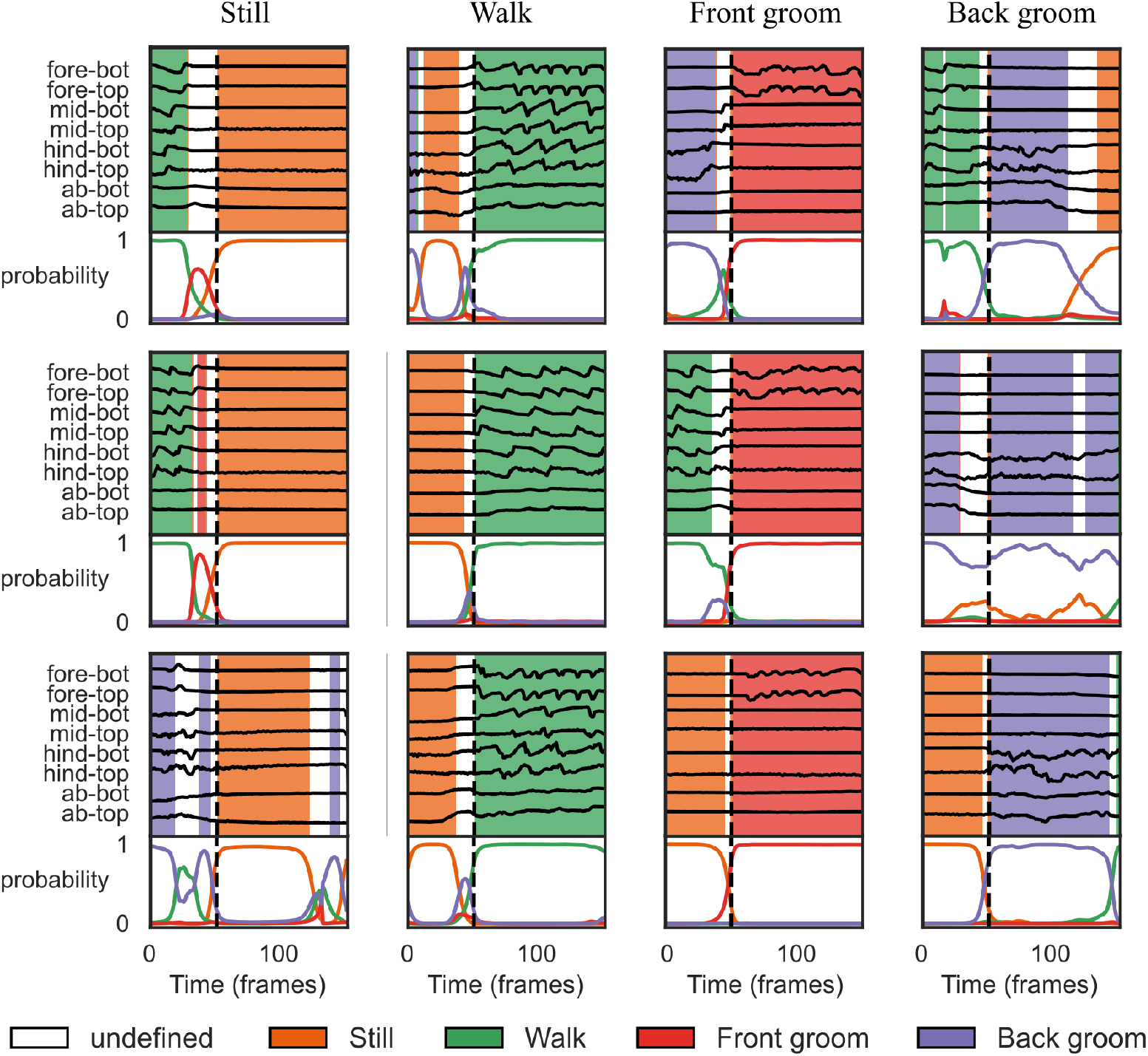
Sequence model outputs for sample time segments. Each panel shows the markers (top, black lines; only x-coordinates are displayed). Background color denotes the highest probability behavior class. The background color is white (“undefined” behavior) if the largest probability is less than an arbitrary threshold of 0.75. The actual class probabilities are plotted below the markers. Each column displays a set of random examples from each behavior class; behavior bout onset is indicated by the vertical black dashed line. Probabilities correspond to the GRU model with *λ*_*h*_ = 0.5, *λ*_*p*_ = 1.0, which achieved the highest F1 score averaged over all behaviors on test data. See Fig. S1 for corresponding outputs of the fully supervised sequence model.

The results above utilize a GRU architecture; to ensure these performance gains are not architecture-dependent, we perform the same hyperparameter search over *λ*_*h*_ and *λ*_*p*_ using two additional architectures: a temporal convolutional network [23, 22] and an MLP neural network with an initial 1D temporal convolutional layer [3, 46] (Fig. S2). We find that for all three architectures the addition of the weak and self-supervised losses drastically improves F1 scores over standard supervised classification with the hand labels.

We next investigate the behavioral embeddings **z**_*t*_ by visualizing them for a single test fly in a 2D space through UMAP [30] (Fig. 4A). The points with corresponding hand labels are colored, revealing that similar behaviors are clustered together. In order to determine how much of this structure can be attributed to the labels, we refit the model with the self-supervised loss only (*λ*_*s*_ = *λ*_*h*_ = 0). The 2D visualization also shows clustered behaviors (Fig. 4B), but with more overlap of different behavior classes. Performing segmentation via clustering on this fully unsupervised behavioral embedding – a standard approach [4, 26, 27] – may therefore result in misclassified behaviors.

**Figure 4.**
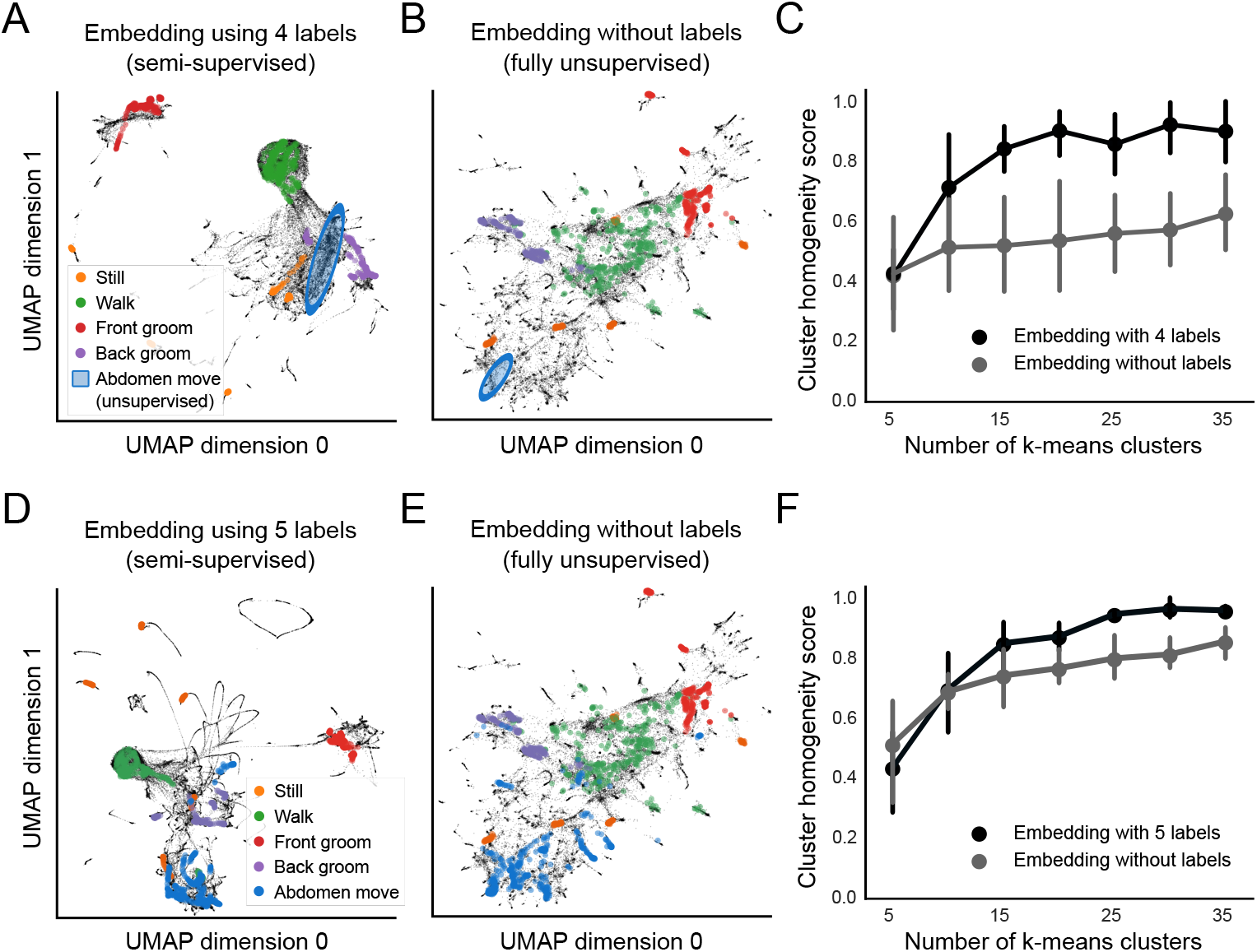
Hand labels improve unsupervised behavioral clustering. *A*: 2D UMAP embedding of behavior colored by hand labels; the model is trained with both hand and heuristic labels from 4 behavior classes: “Still”, “Walk”, “Front groom”, and “Back groom”. *B*: 2D UMAP embedding for the model trained without labels. *C*: The addition of hand labels produces more homogeneous clusters in the 2D space. *D*: Same as panel A, except model is trained with labels from 5 behavior classes - the previous 4 plus “Abdomen move”. *E*: Same as panel B, with additional “Abdomen move” points colored by new hand labels. *F*: Same as panel C, but now cluster homogeneity score is computed with 5 behavior classes.

To quantify this observation, we next perform k-means clustering in this 2D space for both models (with and without labels). We then compute a cluster homogeneity score [37] that measures the extent to which the k-means clusters contain data points from a single behavior class. The model trained with labels achieves a higher score than the fully unsupervised model trained solely with the prediction loss (Fig. 4C). This result demonstrates how adding a small number of hand labels (and/or a larger number of heuristic labels) can improve unsupervised behavioral segmentation with little additional effort.

A primary benefit of unsupervised segmentation is the ability to discover new behaviors in an unbiased manner [9, 35]. For example, one of the k-means clusters from the unsupervised embedding (indicated by the blue ellipse in Fig. 4B) corresponds to periods where the fly moves its abdomen (see marker traces in Fig. S3), a behavior not included in our hand or heuristic labels. Because our semi-supervised model learns to predict behavior at the next time step, it too can potentially capture these unlabeled behaviors. Indeed, we find a k-means cluster in the semi-supervised embedding that also corresponds to this novel abdomen movement behavior (blue ellipse in Fig. 4A).

Next we demonstrate how to utilize this behavioral discovery to further refine the behavioral segmentation. We first hand label the abdomen movement behavior for each of the 10 flies (see Table S1 for details), as well as recompute the heuristic labels: frames are now labeled “Abdomen move” when the ME of the abdomen markers is above a threshold. We retrain the sequence models with these new labels, and find that we can accurately capture this new behavior class (Fig. S3).

We repeat our previous cluster homogeneity analysis, and again find that our semi-supervised approach produces an embedding with a representation of discrete behaviors that is more precise than the fully unsupervised embedding (Fig. 4D-F). Furthermore, we can show that the previous model – trained *without* the “Abdomen move” labels – misclassifies these behavioral bouts as “Back grooming”, “Walk” and “Still” (Fig. 5). Therefore, the addition of this new behavior class refines the previous segmentation.

**Figure 5.**
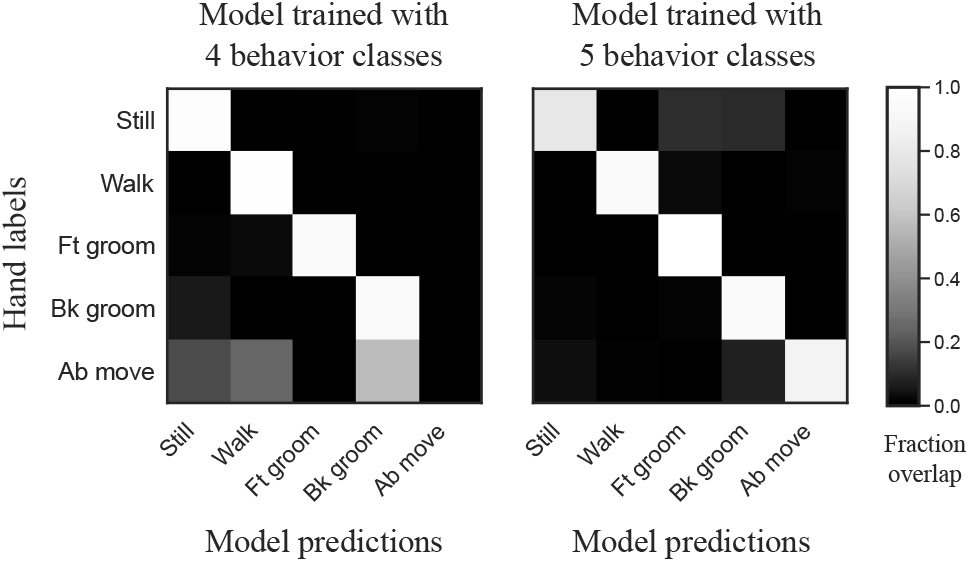
Inclusion of the “Abdomen move” behavior class refines behavioral segmentation. *Left*: Fraction of overlapping frames between the hand labels (y-axis) and model predictions (x-axis) for each behavior class, computed over all test data. Rows sum to 1. Overlap results are shown for best model trained using four behavior classes (same as Figs. 3, 4A, B). *Right*: Overlap results for best model trained using five behavior classes (same as Figs. 4D, E, S3).

## 4. Discussion

We presented an approach to semi-supervised behavioral segmentation that improves upon fully supervised and fully unsupervised approaches. We demonstrated that supervised segmentation metrics (F1) can be improved through the addition of a weakly supervised loss that classifies heuristic labels, as well as a self-supervised loss that predicts the evolution of the behavioral features that serve as model input (Fig. 2). Our work can also be viewed as adding a small number of labels to an unsupervised segmentation problem [10, 25], which we show increases the precision of downstream clustering (thus ensuring the model captures known behaviors of interest) while still allowing the model to discover novel behavioral phenomena (Fig. 4).

This semi-supervised approach can also serve as the foundation for an efficient active learning strategy that reduces human annotation overhead. Computing heuristic labels is a simple strategy to quickly label many frames. A model trained with the weak and self-supervised losses provides an initial embedding that can then guide the selection of frames to label. Additional behavior classes, if discovered, can be added to the set of heuristic and hand labels. This procedure can then be iterated. We demonstrated one aspect of this active learning approach, and believe this is a fruitful direction for future exploration.

## Supplemental Figures and Tables

**Table S1.** The number of labeled frames per behavior for each fly. Initial behaviors (“Walk”, “Still”, “Front groom”, “Back groom”) were labeled in chunks of 50 contiguous time points. Video frame rate is 70 Hz. Flies often engage in behaviors for longer than 50 frames, so the selected 50-frame chunks did not contain any transitions from one behavior to another. The “Abdomen move” behavior was added during a second round of labeling. We labeled longer contiguous chunks in order to capture the full range of the behavior during each bout, which usually consists of raising the abdomen, a brief hold, and then lowering the abdomen. Labeling was performed using the DeepEthogram GUI [6]. The asterisk (**\***) denotes the test experiment visualized in Figs. 3, 4, S1 and S3.

|  | Experiment ID | Total frames | Walk | Still | Front groom | Back groom | Abdomen move |
| --- | --- | --- | --- | --- | --- | --- | --- |
| **Train** | 2019_08_07.fly2 | 50000 | 300 | 300 | 100 | 300 | 591 |
|  | 2019_08_08.fly1 | 94960 | 300 | 300 | 300 | 150 | 594 |
|  | 2019_08_20.fly2 | 65108 | 300 | 300 | 300 | 300 | 1067 |
|  | 2019_10_10.fly3 | 141042 | 300 | 300 | 300 | 300 | 478 |
|  | 2019_10_14.fly3 | 140929 | 300 | 300 | 300 | 300 | 318 |
| **Test** | *2019_06_26.fly2 | 45000 | 300 | 300 | 300 | 300 | 706 |
|  | 2019_08_14.fly1 | 124144 | 300 | 300 | 300 | 300 | 693 |
|  | 2019_08_20.fly3 | 73055 | 300 | 300 | 300 | 300 | 405 |
|  | 2019_10_14.fly2 | 139925 | 300 | 300 | 300 | 300 | 101 |
|  | 2019_10_21.fly1 | 142554 | 300 | 300 | 300 | 300 | 0 |
| **Totals** |  | 1016717 | 3000 | 3000 | 2800 | 2850 | 4953 |

**Figure S1.**
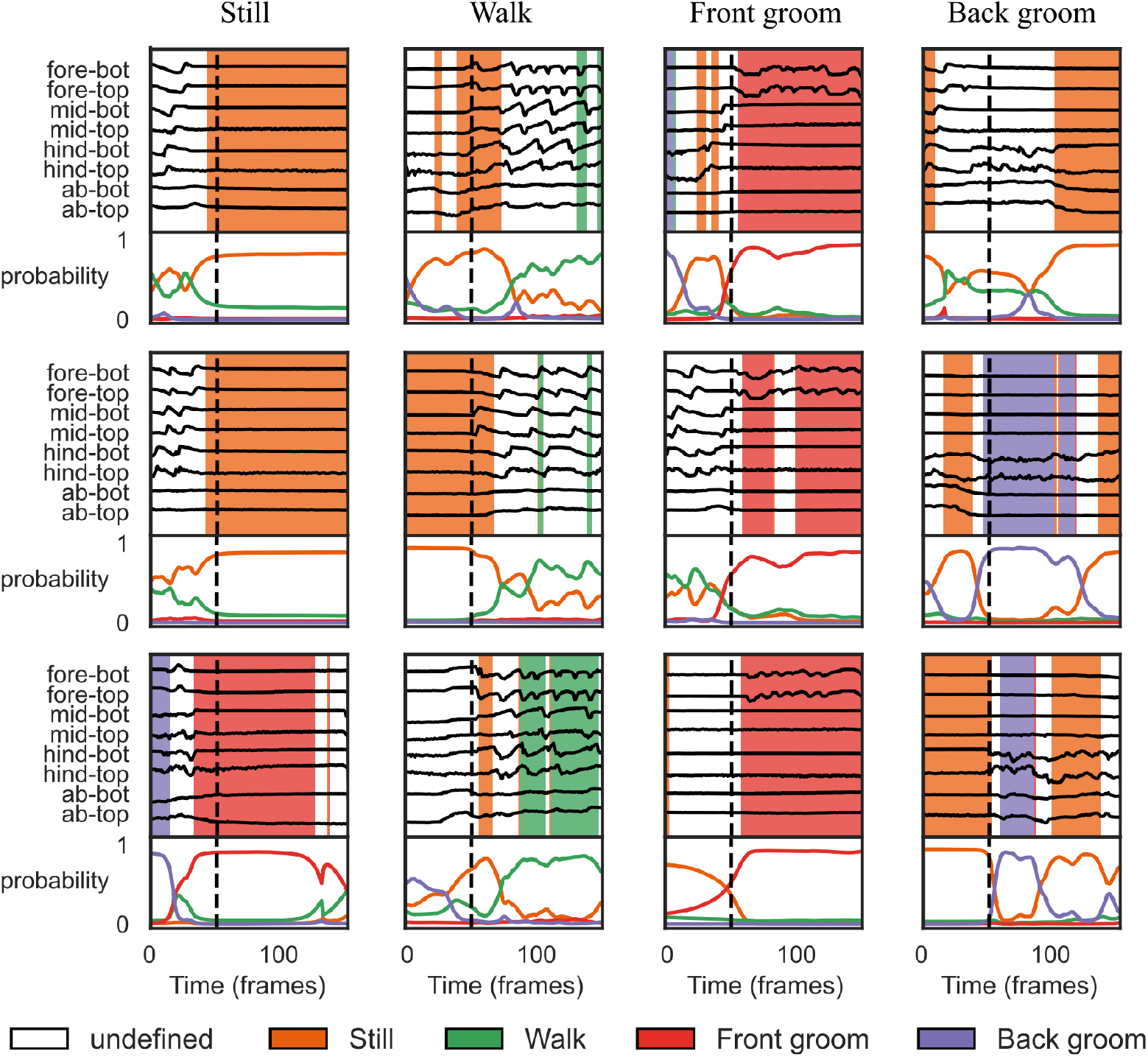
The absence of the weak and self-supervised losses leads to more errors in the segmentation. Panels show the outputs of the fully supervised sequence model (GRU with *λ*_*h*_ = *λ*_*p*_ = 0) for the same sample time segments as in Fig. 3. Note that for some bouts (e.g. those in the “Walk” column) the highest probability belongs to the correct behavior class, but the model is less confident. For other bouts, the model is confident but incorrect (e.g. the final “Still” bout is misclassified as “Front groom”).

**Figure S2.**
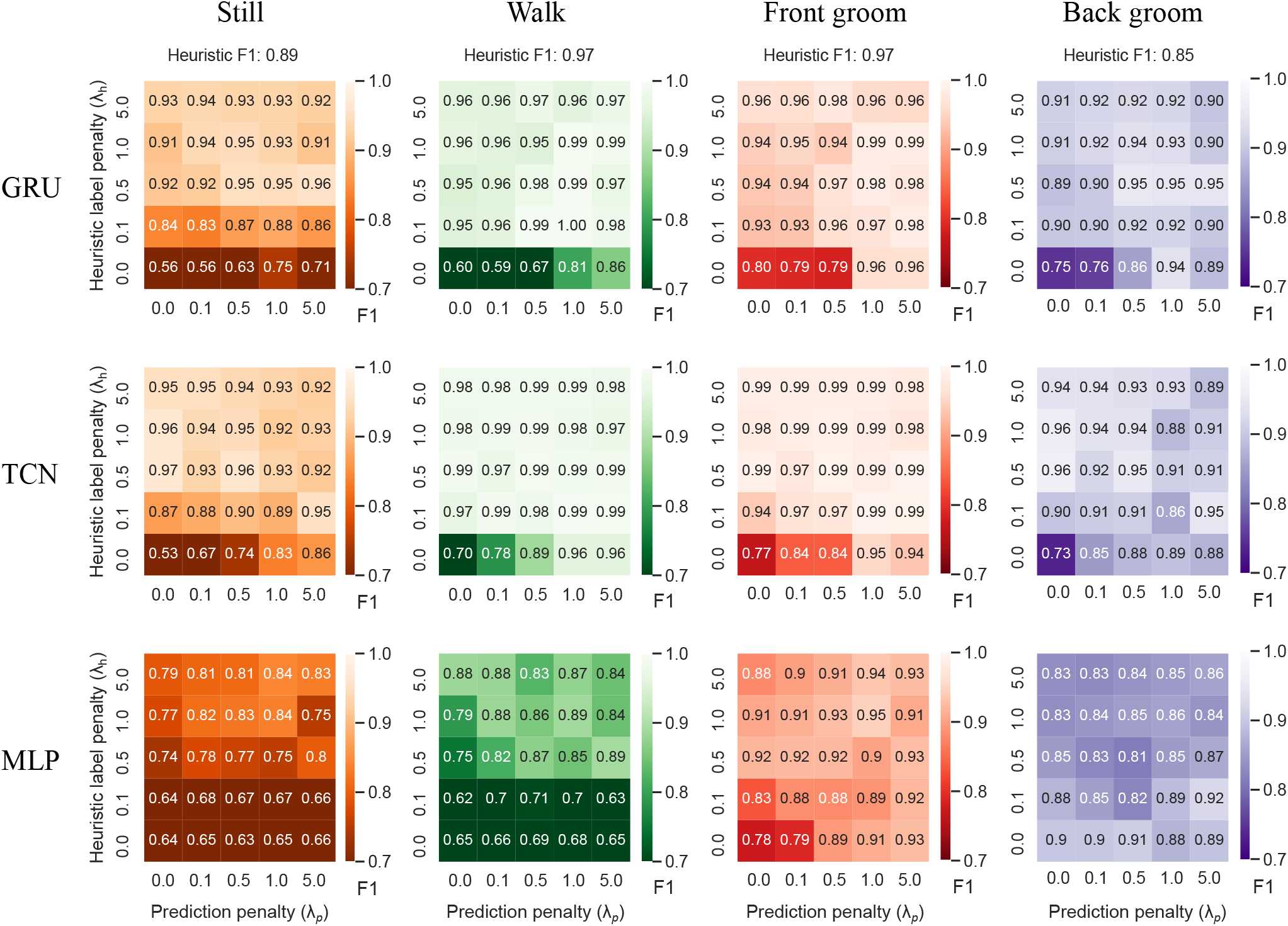
The performance improvement in F1 scores due to weak and self-supervised losses is not architecture dependent. *Top*: F1 results from a GRU where both the encoder and decoder are modeled with two layer bidirectional GRU networks with 32 cells per layer (same models as Fig. 2). *Middle*: F1 results from a temporal convolutional network (TCN) where both the encoder and decoder are modeled with two layers of 1D temporal convolutions (filter size of 17), each followed by a temporal downsampling (encoder) or upsampling (decoder) step. *Bottom*: F1 results from an MLP network where the first layer of the encoder is a 1D temporal convolution (filter size of 17) and the second layer is fully connected (32 hidden units). The decoder is modeled as a two layer MLP (no convolutions) with 32 hidden units per layer. For each architecture we performed a small hyperparameter search across number of layers, cells/units per layer, learning rate, and temporal filter sizes; we found that our results are robust across different settings (data not shown).

**Figure S3.**
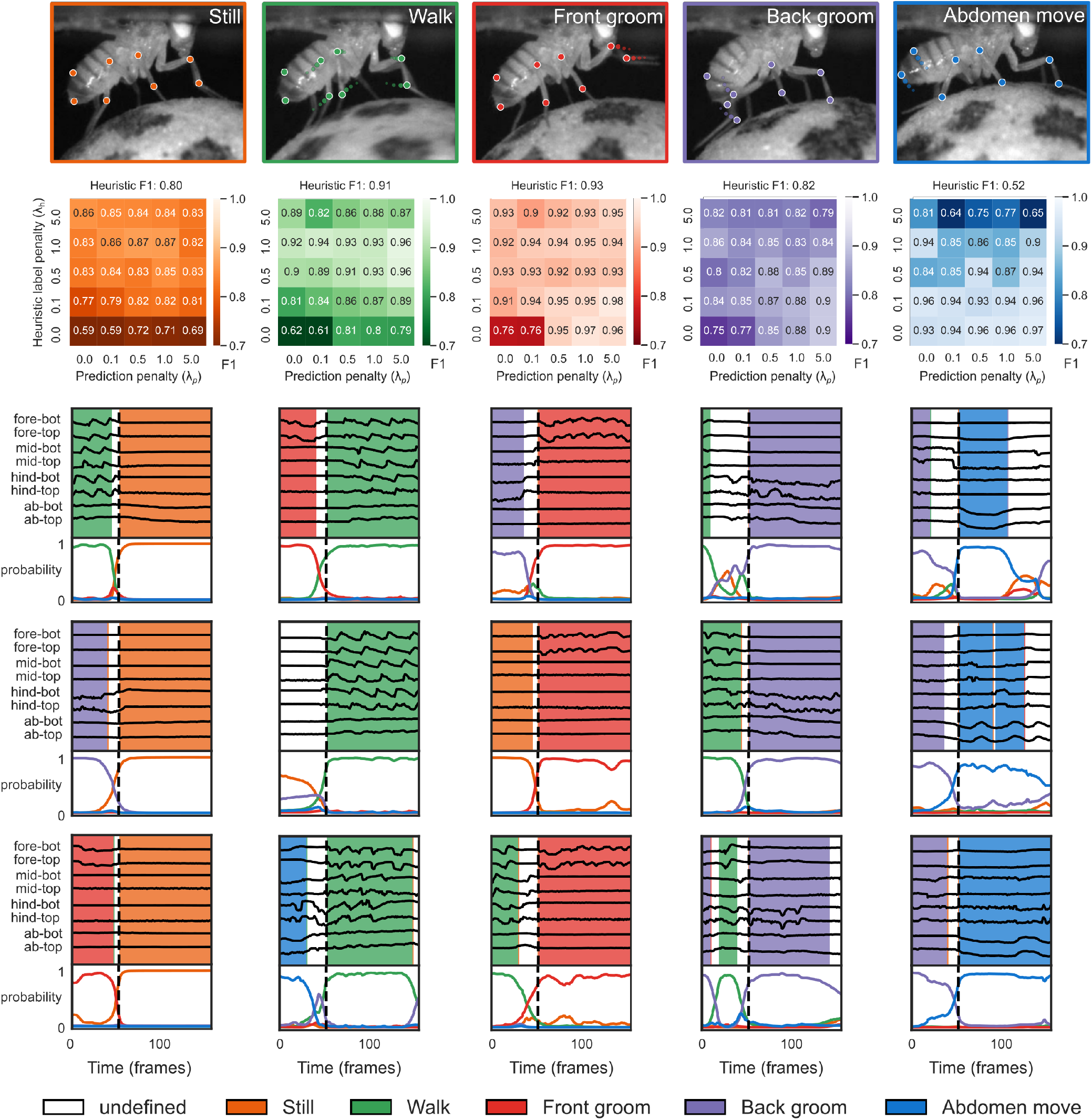
Sequence models can be fit to new behavior classes. Each column represents model fits for a single behavior class. *Top row*: Example frames from five fly behavior classes, overlaid with pose estimates. *Second row*: F1 scores for each behavior on test data. *Bottom rows*: Model outputs for sample time segments. Same conventions as Fig. 3. These probabilities correspond to the GRU model with *λ*_*h*_ = 0.5, *λ*_*p*_ = 5.0, which achieved the highest F1 score averaged over all behaviors on test data.

